# *Thermosipho* spp. immune system differences affect variation in genome size and geographical distributions

**DOI:** 10.1101/331215

**Authors:** Thomas H.A. Haverkamp, Claire Geslin, Julien Lossouarn, Olga A. Podosokorskaya, Ilya Kublanov, Camilla L. Nesbø

## Abstract

*Thermosipho* species inhabit thermal environments such as marine hydrothermal vents, petroleum reservoirs and terrestrial hot springs. A 16S rRNA phylogeny of available *Thermosipho* spp. sequences suggested habitat specialists adapted to living in hydrothermal vents only, and habitat generalists inhabiting oil reservoirs, hydrothermal vents and hotsprings. Comparative genomics of 15 *Thermosipho* genomes separated them into three distinct species with different habitat distributions: the widely distributed *T. africanus* and the more specialized, *T. melanesiensis* and *T. affectus*. Moreover, the species can be differentiated on the basis of genome size, genome content and immune system composition. For instance, the *T. africanus* genomes are largest and contained the most carbohydrate metabolism genes, which could explain why these isolates were obtained from ecologically more divergent habitats. Nonetheless, all the *Thermosipho* genomes, like other Thermotogae genomes, show evidence of genome streamlining. Genome size differences between the species could further be correlated to differences in defense capacities against foreign DNA, which influence recombination via HGT. The smallest genomes are found in *T. affectus* that contain both CRISPR-cas Type I and III systems, but no RM system genes. We suggest that this has caused these genomes to be almost devoid of mobile elements, contrasting the two other species genomes that contain a higher abundance of mobile elements combined with different immune system configurations. Taken together, the comparative genomic analyses of *Thermosipho* spp. revealed genetic variation allowing habitat differentiation within the genus as well as differentiation with respect to invading mobile DNA.

## Introduction

Bacteria of the genus *Thermosipho* are found in high temperature environments such as deep-sea hydrothermal vents and subsurface oil reservoirs. The genus belongs to the phylum Thermotogae and there are currently eight described *Thermosipho* species: *T. africanus, T. activus, T. affectus, T. atlanticus, T. geolei, T. globiformans, T. japonicus* and *T. melanesiensis*. These are thermophilic, organotrophic, sulfur-reducing anaerobes, fermenting various sugars and peptides to H_2_, CO_2_ and acetate (Podosokorskaya et al. 2014).

Comparative genomic analyses of Thermotogae, including *Thermosipho*, have revealed complex evolutionary histories with extensive horizontal gene transfer (HGT), particularly involving members of the bacterial phylum Firmicutes and the Archaea domain (Zhaxybayeva et al. 2009). HGT allows organisms to acquire novel metabolic capacities and adaptations that may lead to species differentiation and innovation (Koonin 2015). In addition, HGT and homologous recombination, may be important in the long-term maintenance and repair of genomic information, by replacing inactivated genes (Takeuchi et al. 2014).

Several observations from *Thermosipho* genomes imply that they are influenced by recent HGT. First, a remnant prophage element was detected in the *Thermosipho melanesiensis* BI429 genome (Zhaxybayeva et al. 2009). Second, in *Thermosipho africanus* TCF52B seventy-eight open-reading frames (ORFs) are annotated as transposases or integrases, and their high intragenomic similarity suggests recent Insertion Sequence (IS) element activity (NesbØ et al. 2009). Both IS elements and prophages are mobile DNA elements and might thus play a role in HGT (Darmon & Leach 2014). And finally, *Thermosipho* genomes carry horizontally acquired genes encoding the vitamin B_12_ synthesis pathway which is absent from other Thermotogae (Swithers et al. 2011).

Intriguingly, thermophile genomes tend to have a higher proportion of HGT acquired genes than mesophile genomes. This might be due to either adaptation to hot environments, DNA repair or dissimilarities of mobile DNA activity rates between hot and mesophilic environments (van Wolferen et al. 2013). Counteracting this trend is the observation that prokaryotic genome size (GS) decreases with an increase of the optimal growth temperature (OGT). This causes thermophiles to have smaller genomes than mesophiles (Sabath et al. 2013; Zhaxybayeva et al. 2012). Sabath et al., (2013) proposed that temperature-induced-genome streamlining was due to a selection for decreased cell and genome size at elevated temperatures. They also showed that many characteristics of streamlined genomes are identified in thermophiles, but it is unclear why thermophiles then have a higher abundance of foreign genes.

Therefore, increasing OGT might increase the selective pressure for mechanisms controlling HGT to prevent unwanted genome expansion. Known vectors of “foreign” DNA, such as phages and transposons, usually have deleterious effects on their host, and most, if not all, prokaryotes have acquired mechanisms that limit their activity. Among these are mechanisms that regulate cellular mobile DNA activity: restriction-modification (RM); abortive infection (ABi); toxin-antitoxin (TA) systems and the clustered regulatory interspaced short palindromic repeats (CRISPR-cas) system (van Houte et al. 2016). CRISPR-cas systems are notably enriched in prokaryotic thermophiles compared to psychrophiles and mesophiles, which implies that HGT control mechanisms are indeed important for thermophile genome maintenance (Koonin et al. 2017).

The understanding of the interaction between GS, HGT and immune system genes is limited for the phylum Thermotogae and the genus *Thermosipho* in particular. For instance, the RM system is well studied and almost universally found in prokaryotes (Roberts et al. 2010), but only poorly characterized among described Thermotogae species (Xu et al. 2011). In contrast, CRISPR-cas systems have been identified in all Thermotogae genomes characterized to date (Zhaxybayeva et al. 2009). Interestingly, CRISPR-cas genes can also be part of mobile DNA itself (Seed et al. 2013), and many show evidence of HGT when comparing for instance *Thermosipho africanus* to *Thermotoga maritima* (Nesbø et al. 2009).

Here we present a comparative analysis of fifteen *Thermosipho* genomes, thirteen of which were generated in this study. The isolates included here were obtained from deep-sea hydrothermal vents and produced fluids from oil reservoirs. The genomes fall into three well-defined lineages (or species) with isolates from the same sample sites showing very high similarities to each other. We show that the three species differ with regard to GS, metabolic repertoire, presence of mobile elements and their immune system gene content, and discuss how the repertoire of immune system genes may influence the observed genomic differences between the three species.

## Results and discussion

### Comparative genomic analysis of Thermosipho isolates

We screened the NCBI-non-redundant (nr)-and IMG 16S rRNA metagenome databases using BLASTn and *Thermosipho* spp. 16S rRNA genes to assess how our genomes span the available environmental diversity of *Thermosipho*. This indicated that most cultivated lineages or species, e.g. *T. melanesiensis, T. affectus* and *T. africanus*, are well covered by our analysis (Figure 1, Supplementary materials Figure S1). Nonetheless, there are several lineages, including *T. geolei, T. ferriphilus* and *T. activus*, for which we do not have genomic data. An IMG database search identified three sequences closely related to *T. africanus* from oil reservoir and hydrothermal vent metagenomes (data not shown). Two recent studies of subsurface crustal fluids also contained *T. africanus* sequences (Nakagawa et al. 2006; Smith et al. 2017), indicating that *T. africanus* is widespread in the subsurface. Moreover, the absence of *T. melanesiensis* and *T. affectus* sequences from other environments than hydrothermal vents, suggests a more specialized lifestyle with limited geographic and ecological ranges.

**Figure 1.**
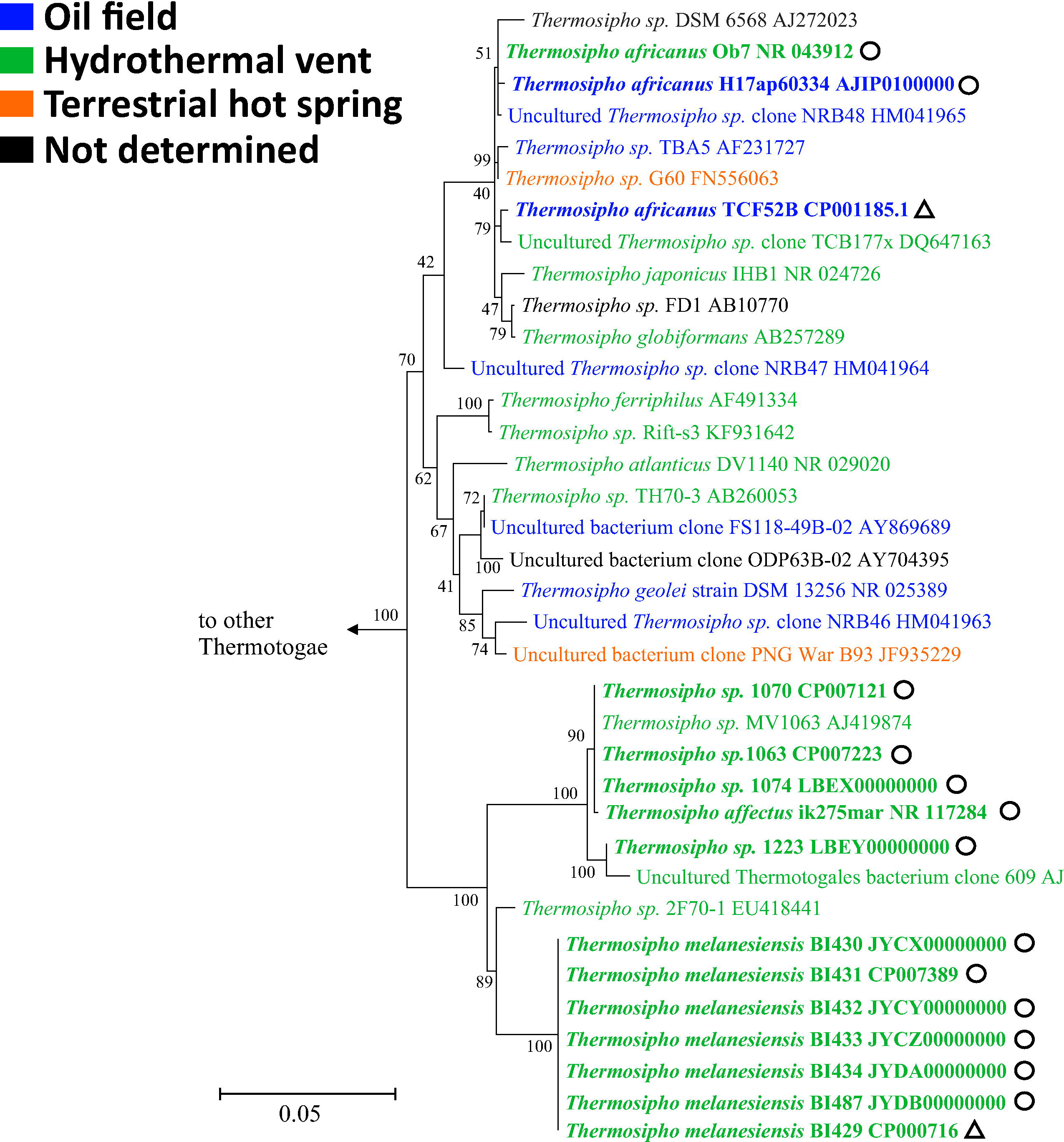
Maximum Likelihood phylogeny of *Thermosipho* 16S rRNA sequences. The tree was constructed in MEGA6 (Tamura et al. 2013) using the General Time Reversible (GTR) model (Γ+I, 4 categories, (Nei & Kumar 2000)). The tree with the highest log likelihood is shown. The percentage of trees in 100 bootstrap replicates in which the associated taxa clustered together is shown next to the branches. Initial tree(s) for the heuristic search were obtained by applying the Neighbor-Joining method to a matrix of pairwise distances estimated using the Maximum Composite Likelihood (MCL) approach. The tree is drawn to scale, with branch lengths measured in the number of substitutions per site, the bar indicates 0.05 substitutions per site. The analysis involved 61 nucleotide sequences. Only alignment positions with fewer than 5% gaps, missing data, and ambiguous bases were included, resulting in a total of 1245 positions in the final dataset. The phylogeny was rooted using 16S rRNA gene sequences of representative sequences from other Thermotogae genera (*Defluviitoga, Fervidobacterium, Geotoga, Kosmotoga, Marinitoga, Mesoaciditoga, Mesotoga, Oceanotoga, Petrotoga, Pseudothermotoga* and *Thermotoga*). All non-*Thermosipho* Thermotogales formed a distinct clade and were collapsed (Supplementary figure S1 for the full phylogeny). *Thermosipho* sequences are colored based on the environment of isolation. Sequences with bold fonts are whole genome sequences. Triangles behind the sequence ID indicate genome not from this study. Circles behind sequence ID indicate genome from this study.

The current study added 13 new *Thermosipho* genomes, with three closed and ten draft genomes, to the two existing ones (Table 1). The overall genomic similarity and phylogenetic relationship of the isolates was calculated by identifying core-genome SNPs (Panseq) (Laing et al. 2010), ANI (Average Nucleotide Identity) (Richter & Rosselló-Mora 2009) and the fraction of shared genes between genomes (Figure 2). These analyses revealed the presence of three distinct lineages. The diversity within each lineage differed and correlated with the geographic range of sampling sites. For example, among *T. melanesiensis* isolates, which were all isolated from the same hydrothermal vent system (Table 1), we found extremely low diversity (intra-cluster ANI values >= 99.98 %), suggesting they are closely related clones. Thus, we kept only the two closed genomes, *T. melanesiensis* BI429 and BI431, in the further analysis. The *T. affectus* genomes, originating from two geographically close sites, show intermediate variation levels. The largest intra-species variation is found among *T. africanus* genomes, which were sampled from three geographically and ecologically separated populations (Table 1). Altogether, these results are consistent with the 16S rRNA findings and suggest that isolation by distance is important for *Thermosipho* spp. differentiation, as observed for other thermophiles (Mino et al. 2017).

**Table 1:**
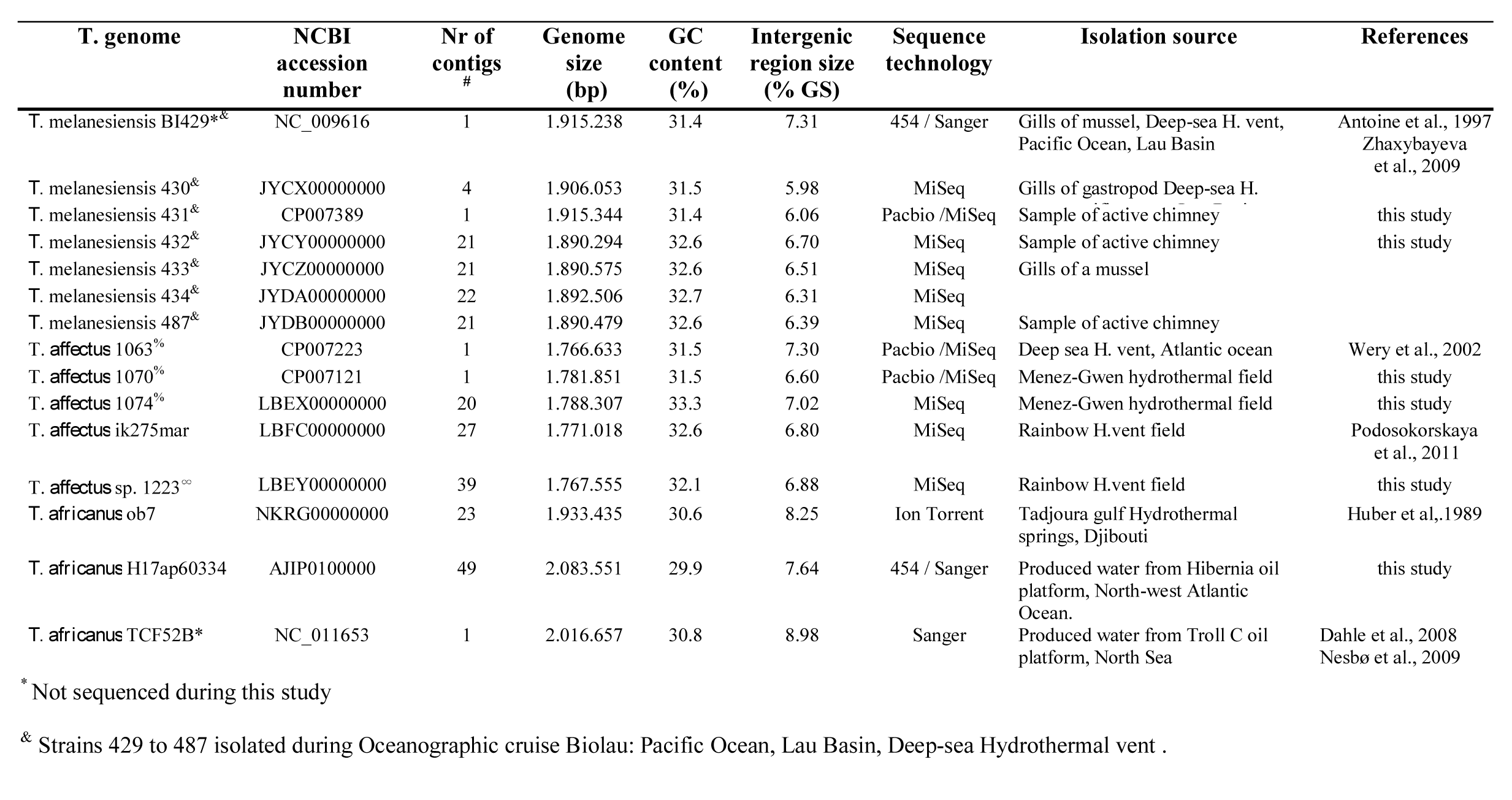
Overview of *Thermosipho* genomes used in this study. % Strains 1063 to 1074 isolated during Oceanographic cruise Marvel: Atlantic ocean, Menez Gwen, Deep-646 sea Hydrothermal vent Isolated during Oceanographic cruise Atos, Atlantic ocean, Rainbow, Deep sea Hydrothermal vent. # Genomes with 1 contig were closed

**Figure 2.**
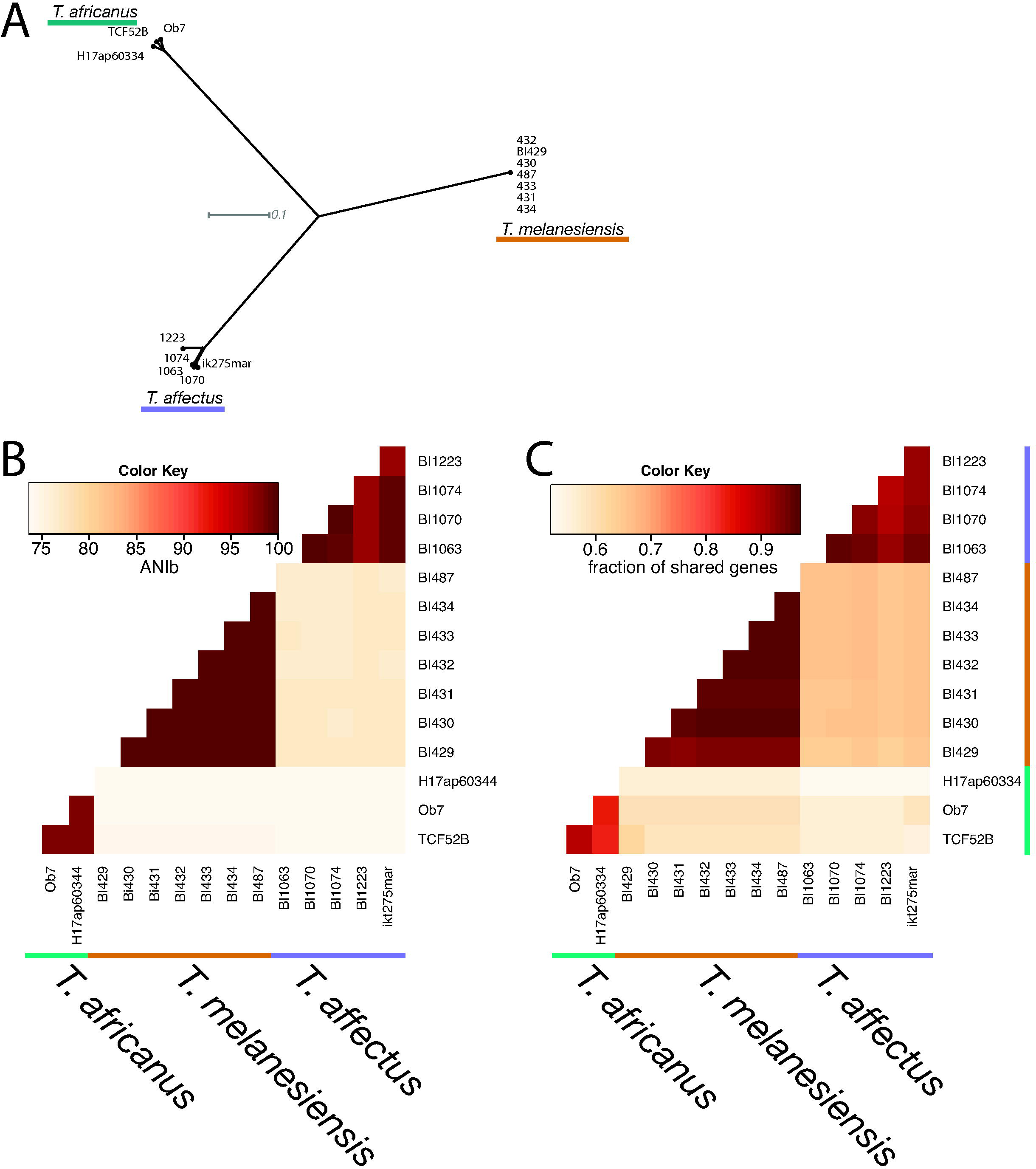
Phylogenomic comparison of 15 *Thermosipho* genomes using core SNPs, ANIb, and the number of shared genes. **A)** Neighbor network of the 15 *Thermosipho* strains. The network is based SNP’s in core genome fragments that were present in all genomes with a minimum of 70% similarity. The network was visualized in Splitstree using the NeighborNet algorithm from uncorrected distances (Huson & Bryant 2006). **B)** Heatmap visualization of pairwise ANIb distances. **C)** Fraction of genes shared between 15 *Thermosipho* genomes. Data was generated at the IMG-database using pairwise Bidirectional Best nSimScan Hits, with genes sharing 70% sequence identity and at least 70% coverage. Percentages were calculated by dividing the number shared genes by the total number of genes for the genome on the y-axis. The colored lines (black, green and blue) indicated which strains belong to which *Thermosipho* lineage/species.

Detection of genomic recombination between prokaryotic isolates can be used to assess species boundaries (Nesbø et al. 2015). LikeWind analysis (Archibald & Roger 2002) detected no recombination events between the lineages suggesting that they indeed correspond to three distinct species (Supplementary materials Figure S2). These results were further supported by pangenome analysis (Figure 2C). Each lineage contained species-specific genes that were at best only very distantly related to genes in the other lineages. These genes mark the *Thermosipho* species boundaries and possibly contribute to niche differentiation between them (Moldovan & Gelfand 2018).

The genomic properties of each strain follow these species boundaries (Table 1; Figure 3A, B). *Thermosipho* genomes have a GC-content (%GC) ranging from 29.9 % (*T. africanus* H17ap60334) to 32.7 % (*T. melanesiensis* 434) with the largest genomes having lowest %GC. GS ranges from 1.77 (*T. affectus* BI1063) to 2.08 Mbp (*T. africanus* H17ap60334), with *T. melanesiensis* strains showing intermediate values (Figure 3A). The *T. affectus* GS places them among the smallest genomes found within the Thermotogae phylum (Figure 3A). Like in other prokaryotic taxa, Thermotogae GS is negatively correlated with OGT, with thermophiles having smaller genomes than mesophilic organisms (Sabath et al. 2013). However, within the *Thermosipho* genus there is a deviation from this trend where *T. africanus* has a larger genome and a higher OGT than the other species (Figure 3A). Additionally, *T. africanus* genomes have more Open Reading Frames (ORFs) compared to the other two species (Table 2) and larger relative intergenic region size (RIRS; 8.29%) compare to *T. melanesiensis* (≈ 6.47%) and *T. affectus* (≈ 6.92 %) (Table 1, Figure 3C). Finally, detection of recently acquired genes showed that the larger *Thermosipho* genomes have more HGT acquired genes than the smaller genomes (Table 2; Supplementary materials Table S1). These results indicate that temperature is not the only factor responsible for the observed *Thermosipho* GS dissimilarities, but that other unidentified processes also influence GS.

**Table 2.** Overview of genome content with a focus on mobile DNA defence systems and mobile elements

| Genome | Closed | ORFs | Species-specific ORFs<sup>#</sup> | Strain specific ORFs | Putative HGT ORFs | Defence genes | CRISPR arrays | Transposase genes<sup>\$</sup> | Prophage present | Prophage ORFs |
| --- | --- | --- | --- | --- | --- | --- | --- | --- | --- | --- |
| *T. melanesiensis* BI429* | Y | 1879 | 424 | 33 | 80 | 34 | 5 | 1 | Yes | 49 |
| *T. melanesiensis* 431 | Y | 1868 | 422 | 4 | 83 | 37 | 5 | 2 (2)<sup>\$</sup> | Yes | 53 |
| *T. sp.* 1063 | Y | 1706 | 349 | 11 | 66 | 31 | 4 | 0 (1) |  |  |
| *T. sp.* 1070 | Y | 1765 | 351 | 43 | 71 | 31 | 4 | 0 (1) |  |  |
| *T. sp.* 1074 |  | 1756 | 353 | 47 | 73 | 31 | 5 | 1 | Yes | 77 |
| *T. affectus* strain ikt275mar |  | 1720 | 348 | 17 | 71 | 33 | 5 | 1(1) |  |  |
| *T. sp.* 1223 |  | 1726 | 349 | 68 | 76 | 30 | 5 | 0 (1) |  |  |
| *T. africanus* ob7 |  | 1902 | 638 | 63 | 83 | 24 | 6 | 10 |  |  |
| *T. africanus* H17ap60334 |  | 1982 | 656 | 169 | 116 | 19 | 6 | 0 (4) | Yes | 48 |
| *T. africanus* TCF52B* | Y | 1954 | 657 | 64 | 77 | 32 | 12 | 43 |  |  |

<sup>#</sup>: Each genome is used as a reference therefore species-specific orfs can show variations.
<sup>\$</sup>: Number in brackets indicates genes with disrupted open reading frames.

**Figure 3.**
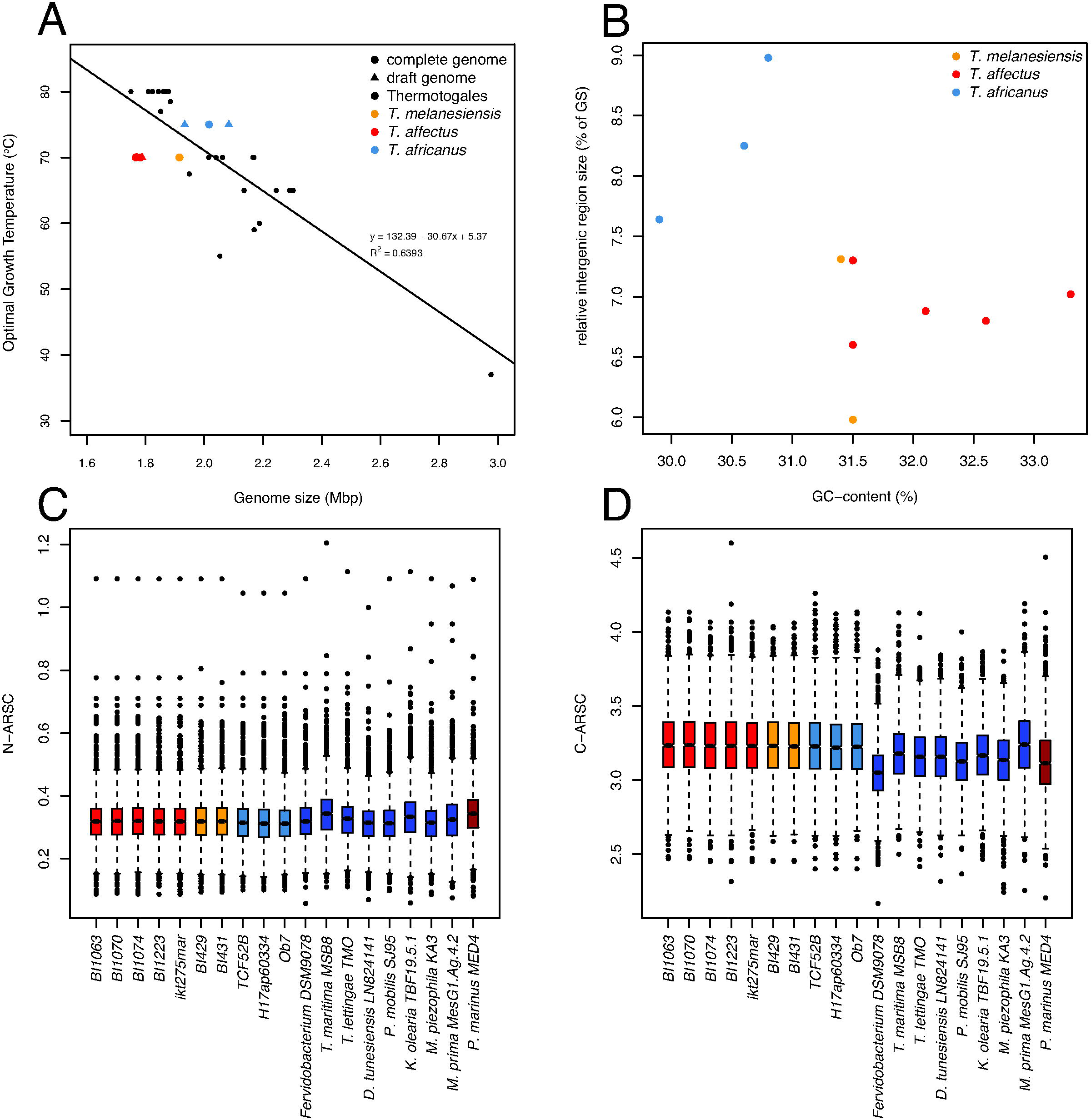
Genomic properties of the *Thermosipho* genomes. **A)** Genome size versus optimal growth temperature for all *Thermosipho* genomes compared to all Thermotogales genomes, for which OGT is known. **B)** %GC versus relative intergenic region size (% of total GS). **C & D) Boxplot** comparison of N-ARSC & C-ARSC values per gene found in *Thermosipho* genomes, one representative genome of each Thermotogales species and *Prochlorococcus marinus* MED4.

Environmental nutrient status was shown to have a positive correlation with prokaryotic GS and %GC, with low nutrient levels selecting for reduced GS and %GC via genome streamlining or by increasing the mutational load (Mende et al. 2017; Gupta et al. 2016; Giovannoni et al. 2014). Thus, under oligotrophic conditions prokaryotes experience selective pressure against the usage of high nitrogen content amino acids (aa). Accordingly, preference for aa with lower nitrogen and higher carbon content could be an indicator for nitrogen limited environments (Grzymski & Dussaq 2011; Mende et al. 2017). An analysis of average nitrogen and carbon molecules per aa-residue-side-chain (N-ARSC and C-ARSC) showed that *Thermosipho* genes have a median N-ARSC of 0.32 and a C-ARSC of 3.2 (Figure 3C, D). Such N/C-ARSC values are similar to what is found in other Thermotogae and also in streamlined bacteria living in oligothrophic environments (e.g. *Prochlorococcus marinus* strain MED4) (Figure 3C, D)(Grzymski & Dussaq 2011), and are consistent with Thermotogae lineages being members of the nutrient poor and nitrogen limited subsurface biosphere (Head et al. 2014). These results could explain the generally small Thermotogae genome sizes.

A characteristic of small genomes is the presence of few paralogous genes. When considering all *Thermosipho* genomes, on average 14.7% of the COGs have paralogs while 85.3% (+/-0.7 %) of the COGs are single-copy genes. These numbers are similar to those of other streamlined genomes (Grote et al. 2012). A comparison of COG annotations (Figure 4A; Supplementary Materials Table S2) revealed several categories with significant differences in relative abundance between the three species (p ≤ 0.01 for B, C, H, I, J, O and R and p ≤ 0.05 for G and L). Two COG categories show large relative abundance differences between species that together may explain a significant fraction of the GS differences.

**Figure 4.**
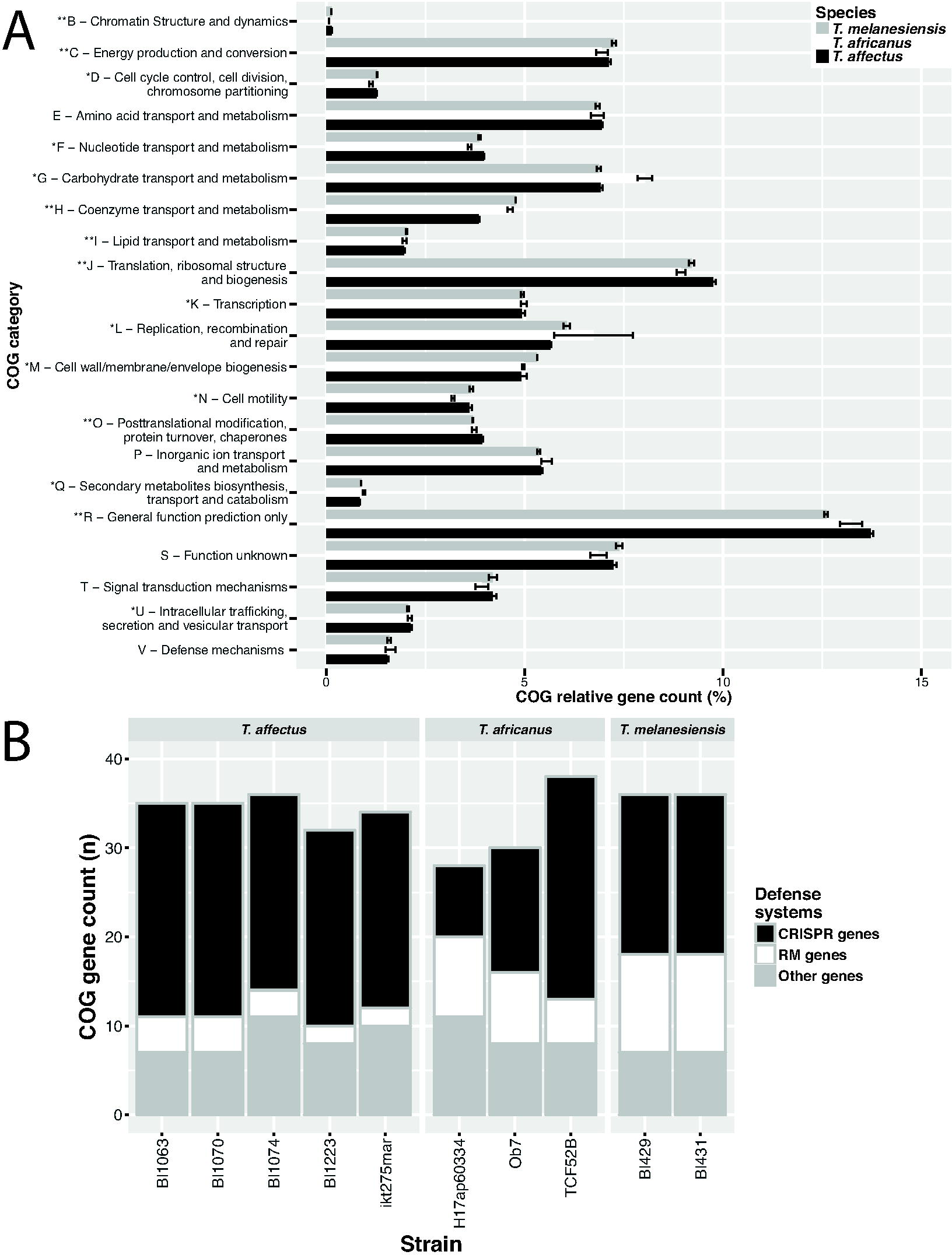
Comparison of COG functional annotations for three *Thermosipho* species. **A)** Complete genome COG category annotations averaged for three *Thermosipo* species. COG category counts are relative to indicate proportional differences in gene content between the three species. The three species are indicated in the textbox in the figure. **B)** Mobile DNA defense related COG annotation counts for 10 *Thermosipho* isolates. A total of 38 COGs were identified using a list of COGs related to defense genes (Makarova et al. 2011). The counts of the identified COGs were summarized into three groups: Restriction-modification systems COGs, CRISPR-cas associated COGs, and other COGs. The strains are grouped per *Thermosipho* species.

For the first category, COG G (Carbohydrate transport and metabolism), *T. africanus* genomes have more genes than the other two species (Figure 4A). Similarly, PFAM based comparisons using the carbohydrate database dbCAN (Yin et al. 2012) (Supplementary information; Supplementary materials Table S3), identified several enzyme families involved in breakdown of various beta-linked oligo-and polysaccharides and some of these were exclusively found among *T. africanus* representatives. Taken together with phenotypic differences (Podosokorskaya et al. 2014), this suggests that *T. africanus* spp. might be more versatile with regard to carbohydrate uptake and metabolism than the other two species and this could be at least partly responsible for its wider ecological distribution.

For the second category, COG H (coenzyme transport and metabolism), the large relative abundance difference is due to *T. affectus* genomes lacking most (20 out of 22) genes needed for corrinoid synthesis. Corrinoid synthesis is an essential part of *de novo* vitamin B_12_ synthesis and the lack of these genes suggests that *T. affectus* only rely on vitamin B12 precursor molecule recovery (Supplementary materials Figure S3). In contrast, a complete set of corrinoid synthesis genes is found in *T. africanus* and *T. melanesiensis* genomes, which are able to synthesize vitamin B_12_ (Swithers et al. 2011) (Supplementary materials Figure S3). It is unclear how this vitamin B_12_ pathway difference affects *Thermosipho* spp. ecology, but the loss of part of this system could be interpreted as a way to reduce metabolic costs under nutrient limited conditions.

GS in prokaryotes can be influenced by the presence of genomic islands, which often encode IS elements or prophages (Hacker & Kaper 2000). The *T. africanus* genomes have the highest abundance of transposase genes, while the two other species have few or only disrupted transposases (Table 2). Besides IS elements each isolate contains blocks of co-localized genes unique for that genome (cluster size range: 4 – 31 genes). Interestingly, several of these regions encode likely integrated prophages (*T. melanesiensis* BI429 / BI431, *T. affectus* BI1074, *T. africanus* H17ap60334) as they contain typical phage genes (Supplementary information, Supplementary materials Table S4). Recent reports have identified three temperate siphoviruses in Thermotogae genomes, and together with the above findings, it becomes clear that prophages are not unusual genomic features within this phylum (Lossouarn et al. 2015; Mercier et al. 2018). The remaining clusters are dominated by hypothetical proteins, CRISPR-cas genes or genes involved in various types of cellular activities (Supplementary materials Table S4). These findings indicate higher overall mobile element abundance in *T. africanus* genomes compare to both other species, which partially explains the larger GS of this lineage.

### *The* Thermosipho *spp. immune systems*

The mobile element differences among *Thermosipho* spp. suggest dissimilarities between species in their foreign-DNA defense mechanisms. The *Thermosipho* genomes contain 44 COGs involved in such defense systems (Makarova et al. 2011); RM system (12), CRISPR-cas genes (18) and genes with other COG annotations (14) (Figure 4 B).

The function of CRISPR-cas genes is to recognize and degrade mobile elements using CRISPR spacers. Each genome contains mostly unique spacers with only a few shared within species (Supplementary materials Figure S4). Considering CRISPR-cas genes, most are found in *T. affectus* isolates and *T. africanus* TCF52B, while *T. africanus* Ob7 and H17ap60334 genomes have the least. All genomes contain CRISPR-cas Type III systems (A & B, targeting RNA) while some have also Type I (targeting DNA) (Tamulaitis et al. 2017). The Type III system requires *cas10* (COG1353) and is detected at multiple loci in all genomes, except for *T. africanus* that have one gene (Type III-B). CRISPR-cas Type I requires *cas3* (COG1203) (Makarova et al. 2015). This gene is present in *T. affectus* BI1223 and *T. africanus* TCF52B, but disrupted in *T. melanesiensis, T. africanus* Ob7, and absent in *T. africanus* H17ap60334 (Supplementary materials Table S5 / S6). Finally, the remaining *T. affectus* genomes have a divergent *cas3* (NOG212552) (Supplementary materials Table S5 / S6). Overall, we found that CRISPR-cas systems differ across the genomes, which likely results in different CRISPR-cas immune responses among *Thermosipho* spp.

The RM system utilizes genome methylation and restriction enzymes to identify and attack foreign DNA. *T. affectus* genomes lack restriction enzymes, suggesting the absence of a functional RM system (Figure 4B). Pacbio sequencing of two *T. affectus* isolates (BI1063 / BI1070) indicated absence of genome methylation. In contrast, a methylated genome was detected in *T. melanesiensis* BI431 together with type I, II and III restriction enzymes and DNA methylases. All the *T. africanus* genomes have a complete type III system, while type II is only present in strains Ob7 and H17ap60334 (Supplementary materials Table S5 / S6). It is unclear if these genomes are methylated, since Pacbio sequencing was not performed.

The above analysis shows differences in foreign-DNA defenses between three *Thermosipho* species. Since each defense system can impose different fitness cost on the host (van Houte et al. 2016), our results suggest that each *Thermosipho* spp. experiences different phage predation pressure. For instance, when phage diversity is high, a non-specific defense system like the RM system might give a selective advantage despite increased fitness costs due to autoimmune reactions affecting host genome integrity. In contrast, the CRISPR-cas system is an inducible and highly precise response offering resistance to specific phages, and is optimal at low phage diversity and under small infection risk. Such immune systems differences may lead to long-term evolutionary implications for acquiring novel DNA via HGT or recombination. Several studies showed that: i) organisms that rely on HGT lack CRISPR-cas systems since it can block HGT; ii) genomes without CRISPR-cas tend to be larger than closely related isolates with CRISPR-cas present, and iii) phase-variation of RM systems, switching of gene activity and phenotype, can lead to uptake of beneficial DNA which allows for adaptive speciation (reviewed in (van Houte et al. 2016)).

Therefore, based on these fitness effects of their immune systems, we propose the following interpretation of the observed GS differences. The presence of different CRISPR-cas types in *T. affectus* genomes has likely limited mobile element occurrence in those isolates. In addition, these systems may have induced genetic isolation and speciation by also limiting DNA recombination. In *T. melanesiensis* we find a complete CRISPR-cas Type III system and a full scale RM-system. This is able to tolerate the presence of a prophage in the genome, which would likely not be possible if an active Type I system was targeting the specific phage (Pyenson & Marraffini 2017). In contrast, the *T. africanus* defense systems with a more reduced CRISPR-cas and RM systems set-up, may have allowed genomic expansion by increased recombination via HGT and by higher mobile element activity. This could further have induced, or maintained, a larger metabolic potential as observed from the carbohydrate genes, permitting this species to thrive in a wider variety of ecosystems than the metabolically less diverse *T. melanesiensis* and *T. affectus* species.

## Conclusions

Here we described three clearly separated species-level genomic lineages within the genus *Thermosipho*: *T. affectus, T. africanus* and *T. melanesiensis*. Recombination detection showed very little interspecies but high intraspecies gene flow. This finding was supported by differences in GS, RIRS, %GC, and the presence of large species-specific gene sets, reflecting the metabolic differences between species. The species-specific genes sets probably influence distribution differences of *Thermosipho* species in various subsurface ecosystems. *Thermosipho* bacteria frequently interacted with mobile DNA such as phages and transposons that were identified in several genomes. The differences in abundance of mobile elements among the three species are likely caused by immune system differences. Interestingly, the nature of the immune system could play a large role in genome maintenance and ecological adaptation, by differential limitation of genomic recombination via HGT. Taken together, our results suggests that genomic and phenotypic differences between the three *Thermosipho* species can be interpreted as the combinatorial result between genome streamlining and immune system composition.

## Materials & Methods

### DNA extraction and genome sequencing

Thirteen *Thermosipho* strains were cultured in a modified Ravot medium as previously described in (Lossouarn et al. 2015) and used for total DNA extraction as described in (Charbonnier et al. 1992; Geslin et al. 2003). All strains, except *T. africanus* Ob7 were submitted for genome sequencing at the Norwegian Sequencing Centre (NSC, Oslo, Norway). A 300 bp insert paired end library was prepared using the genomic DNA sample prep kit (Illumina, San Diego, CA, USA) for each DNA sample. The Library was spiked with phiX DNA. Subsequently, all strains except Ob7 were sequenced on an Illumina MiSeq generating a dataset of 250 bp paired end reads.

Strain Ob7 was sequenced at University of Alberta, Canada, using an Iontorrent (Thermo Fisher Scientific, Waltham, MA, USA) approach. DNA was enzymatically sheared using the Ion Shear Plus kit (Life Technologies, Carlsbad, CA, USA) then cloned using the Ion Plus Fragment Library kit (Life Technologies) following the manufacturer’s instructions. The library was then sequenced on an Ion Torrent PGM using a 316D chip and 500 flows.

Four isolates (*Thermosipho* sp. 430, 431, 1063, 1070) were selected for long read sequencing with the goal to complete the genomes and assess methylation status of the DNA. DNA’s were prepared following the Pacific Biosciences instructions and sequenced at the NSC on a Pacbio RSII (Pacific Biosciences, Menlo Park, CA, USA).

### Quality control and genome assembly

Pacbio sequencing of the strains 431, 1063 and 1070 produced libraries suitable for genome assembly without the addition of Illumina paired-end reads. For all strains Pacbio read filtering and error-correction were performed using version 2.1 of the Pacbio smrtportal (http://www.pacb.com/devnet/). Assembly and base modification detection were performed using the RS_HGAP_Assembly.2 and RS_Modification_and_Motif_analysis protocols.

Overlapping regions between contigs were manually detected using a combination of read mapping and subsequent visualization with IGV (v. 2.3.23) (Thorvaldsdottir et al. 2013). This allowed us to inspect regions with low quality mappings (due to the presence of duplicated regions), and similar gene annotations on both sides of a contig gap. Preliminary annotations were prepared using Glimmer at the RAST server and inspected in the CLC Bio main workbench (Overbeek et al. 2013). Bam files were generated by mapping MiSeq PE reads using bwa version (v. 0.7.5a) to the annotated contigs (Li & Durbin 2009) and by converting sam files with Samtools (v. 0.1.19). Bam files were imported into the IGV viewer together with the annotated contigs. This approach resulted in closed chromosomal sequences.

For strain 430, we generated a combined assembly using high quality illumina MiSeq reads and a smaller set of Pacbio reads (15594 subreads) in an assembly using Spades (v. 3.5.0.) (Bankevich et al. 2012) with kmer size set to 127. Low coverage contigs (< 2.0) were discarded. The remaining contigs were checked in IGV for miss-assemblies using mapped MiSeq and Pacbio reads and if detected contigs were discarded. Finally the contigs were checked for unambiguously overlapping ends as described above and if detected the contigs were combined.

The *T. africanus* Ob7 genome was assembled using NEWBLER v. 2.6 (Margulies et al. 2005). The remaining strains, were assembled using CLC assembler v.7, Spades (v. 3.5.0.) and Velvet (v 1.2.10) (Zerbino & Birney 2008) (Table 1). Genome assemblies were compared using REAPR (v1.0.16) and the most optimal assembly for each strain was selected for genome annotation with the PGAP pipeline from NCBI (Hunt et al. 2013).

### 16S rRNA Phylogeny of Thermosipho species

Full-length 16S rRNA sequences were manually extracted from the annotated *Thermosipho* genome assemblies generated in this study as well as publically available genomes. Next we searched the SILVA database version SSU r122 for *Thermosipho* spp. 16S rRNA sequences with the following setting: sequence length > 1300 bp, alignment quality > 80, pintail quality > 80 (Quast et al. 2013). Matching sequences were downloaded and added to extracted 16S rRNA sequences from the de-novo sequenced genomes. Multiple identical sequences from the same strain with different accession numbers (e.g. the four 16S rRNA sequences of *T. melanesiensis* BI429, five 16S rRNA sequences *T. africanus* TCF52B) are represented by a single sequence in the dataset.

The NCBI non-redundant database was then queried using BLASTn (blast+ v 2.2.26 (Camacho et al. 2009)) with the 16S rRNA gene dataset described above, to identify *Thermosipho* sp. like 16S rRNA sequences missed with the above methods. Additional full-length sequences were downloaded from Genbank and combined with the de-novo / SILVA 16S rRNA sequences.

The final 16S rRNA gene alignment consisted of *Thermosipho* sequences combined with outgroup sequences from other Thermotogales species. The resulting alignment was used to build phylogenies using Maximum Likelihood as implemented in MEGA 6.0 (Tamura et al. 2013) and the General Time Reversible (GTR) model (G+I, 4 categories) as suggested by JModelTest v. 2.1.7 (Darriba et al. 2012).

### Genome properties and selection analysis

The %GC of the three positions in each codon of all genes was using the software codonW version 1.4.4 (http://codonw.sourceforge.net/). The N/C-ARSC values were calculated with the python scripts from the publication (Mende et al. 2017) after modifying them for usage on our genomes.

### Pangenome analysis

Blast Ring genome plots were generated using BRIG version 0.95 (Alikhan et al. 2011) running Blast+ version 2.2.28 (Camacho et al. 2009) with the following set up. The nucleotide sequence of one of the fifteen *Thermosipho* genomes was used to create the blast-database with default settings. Nucleotide sequences of all coding genes were extracted from each genome (NCBI Genbank annotation) using CLC Main workbench Version 6.8.3 and were used for BLASTn analysis in BRIG with the following settings: max_target_seqs: 1; max e-value cut-off: 1.0^-4^. Alignments with a minimum of 70% similarity were visualized with BRIG. The same procedure was used for all genomes.

Complete chromosome sequences or contigs were uploaded for all fifteen genomes to the Panseq 2.0 server (https://lfz.corefacility.ca/panseq/) for pangenome analysis (Laing et al. 2010). Sequence similarity cut-off (SC) was set at 70 % to identify core-genome segments and SNP’s. We used the standard settings with Panseq except: percent sequence identity cut-off set to 70%, core genome threshold: 15, BLAST word size: 11. The final alignment of the single nucleotide polymorphisms (SNPs) was loaded into splitstree and visualized as an unrooted phylogeny using the neighbor network algorithm (Huson & Bryant 2006)

Ten genome sequences (none-closed genomes of the *T. melanesiensis* cluster were excluded, due to high sequence similarity) were aligned with progressiveMauve (Mauve (v. 2.3.1) (Darling *et al.*, 2010)) using *T. africanus* H17ap60344 set as the reference and automatically calculated seed weights and minimum Locally Colinear Blocks (LCB) scores. Gaps were removed and the edited LCBs were concatenated in Geneious 8 (www.geneious.com).

Recombination analysis of the concatenated alignment was done in LikeWind (Archibald & Roger 2002) using the maximum likelihood tree calculated in PAUP* version 4.0b10 (Swofford 2002) under a GTR+Γ+I model as the reference tree.

Pairwise Average Nucleotide Identity Blast scores (ANIb) were calculated using the JspeciesWS webtool (http://jspecies.ribohost.com/jspeciesws/) (Richter & Rosselló-Mora 2009; Goris et al. 2007). The results were visualized with R (version 3.2.1) using the heatmap.2 function (Package gplots) and the *Thermosipho* genomes were clustered using Jaccard distances based on pairwise ANIb values (Package Vegan. version 2.3-0).

The IMG/ER Pairwise ANI calculation was used to determine the number of shared genes between each genome separately (Markowitz et al. 2009). The method uses pairwise bidirectional best nSimScan hits where similar genes share a minimum of 70% sequence identity with 70% coverage of the smaller gene. For each genome we calculated the fraction of shared genes with all other genomes. Unique genes per genome were determined using the IMG Phylogenetic profiler tool for single genes, where each genome was analyzed for the presence of genes without any homologs in all other 14 genomes. The settings were: minimum e-value of 1.0^-5^; min percent identity: 70%; pseudogenes were excluded; the algorithm was present/absent homologs. The same tool was used to identify the presence of homologs shared with all genomes, and with only the genomes from the same cluster.

### Functional comparison of genome

The fifteen genomes were compared using the Clusters of Orthologous Genes (COGs) annotations. A COG reference database (version 10) was downloaded from the STRING database (http://string-db.org/). For each genome all protein sequences were aligned using BlastP, with the settings: 1 target sequence; maximum e-value: 1.0e^-20^; database size 1.0^7^; tabular output. Only hits to COG database sequences were retained when the alignment was >= 70% of the length of the longest protein sequence and were used to build a protein COG classification table. The COG IDs were summarized by classifying them to any of the available COG categories (ftp://ftp.ncbi.nih.gov/pub/wolf/COGs/COG0303/cogs.csv). For the COG analysis of the species-specific genes, we used the results from the IMG phylogenetic profiler tool to identify cluster specific genes. The COG classification table was screened for cluster specific genes and summarized into COG categories. Next, we searched for paralogous genes by examining how many orthologous genes, classified as COGs, were assigned per COG. This identified COGs only present once or multiple times in the genomes, and those results were summarized.

In order to statistically compare species differences of COG categories we normalized counts by total COG gene annotations, giving relative abundances per genome. R (version 3.3.0) was used to identify COG categories that were significantly different between the three species using the non-parametric Kruskal-Wallis test (p-value <= 0.01). The R-package ggplot2 (version 2.1.0) was used to generate the comparisons graphically.

### Horizontal gene transfer detection

Genes putatively acquired by HGT were identified using HGTector (Zhu et al. 2014) with BLASTp (blast+ v 2.2.26). The databaser.py script was used on 6 December 2015 to download per species one representative proteome of all microorganisms from the NCBI refseq database. We compared the predicted *Thermosipho* spp. protein sequences to the reference database using the following BLAST cut-offs: E-value: 1.0^-5^, Percentage identity: 70%, percentage coverage: 50%, and a maximum of 100 hits were returned. To determine which genes were putatively acquired by any of the strains we set the HGTector self group to the genus *Thermosipho* (NCBI taxonomy ID: 2420) and the close group to either the family Fervidobacteriaceae (NCBI taxonomy ID: 1643950), or the order Thermotogales (NCBI taxonomy ID: 2419). HGTector then analyzes the blast output from each protein for hits matching taxa belonging to either the self or close groups, or more distantly related taxa, which is used to determine which genes have likely been acquired from taxa more distantly related than the close group. When the close group was set to Fervidobacteriaceae, we identified putative HGT genes specific for *Thermosipo* sp. but not found in Fervidobacteriaceae species. This setting however, did not indicate putative HGT genes derived from other families within the Thermotogales order or beyond.

### Defense genes

Using the defense genes list created by Makarova et al., (2011), we screened the IMG genome Clusters of Orthologous Genes (COG) annotations for the presence of any COGs involved in any of the mobile DNA defences (Makarova et al. 2011). We summarized the identified COGs and distinguished between CRISPR-cas associated or restriction-modification (RM) system genes. CRISPR-cas genes were manually inspected to determine completeness / disruption of the genes. In addition, did we manually inspect IMG gene annotations for the presence of CRISPR-cas genes not identified with the Makarova et al., (2011) database. This was used to identify the CRISPR-cas gene clusters and determine the CRISPR type encoded by the genes in a cluster.

### Crispr-spacer analysis

All genomes were uploaded to CRISPR finder (http://crispr.i2bc.paris-saclay.fr/) to detect CRISPR-arrays (Grissa et al. 2007). CRISPR spacer sequences from each genome were compared using BLASTn against all *Thermosipho* sp. spacer sequences using the following settings: e-value cut-off: 1.0^-5^, database size: 1.0^7^, dust: no. The tabular blast results were visualized with R-statistics using the Markov Cluster Algorithm (MCL) (v1.0) and Igraph (v1.0.1) packages. Igraph was used for matrix construction. MCL was run using the matrix with the inflation set to 1.4 and max iterations set to 100 (Enright et al. 2002).

In order to search for matching sequences within the genome but outside the CRISPR arrays, e.g target genes, we masked all CRISPR arrays using maskfeat (EMBOSS v. 6.5.7 (Rice et al. 2000)). Next we ran BLASTn (v2.2.26+) using the spacers of each genome against the own genome using the following settings: e-value cut-off: 1.0^-5^, database size: 1.0^7^, dust: no. CRISPR array spacers were also compared against the NCBI nucleotide database to find other species with similar sequences. BLASTn was run with the settings: e-value cut-off: 1.0^-5^, dust: no. Each genome was screened for the presence of prophages (Supplementary information for details).

### Vitamine B_*12*_ pathway analysis

The genes involved in the Vitamine B_12_ metabolism are found in four different gene clusters (BtuFCD, Corriniod, Cobalamin, and SucCoA) in *Thermosipho* and can be regulated by B_12_ riboswitches (Swithers et al. 2011). All 15 genomes were screened for the presence of Cobalamin specific riboswitches using Riboswitch scanner (Mukherjee & Sengupta 2015). This information was used to confirm the presence of the four gene clusters in each genome. Next, we extracted the protein sequences from the *T. melanesiensis* BI429 genome involved in B_12_ metabolism (Swithers et al. 2011) and used them to identify homologous genes in all *Thermosipho* genomes using tBLASTn with a maximum e-value 1.0^-20^.

### Data desposition

All genomes were deposited in the Genbank database and their accession numbers are found in Table 1. In addition all genomes were deposited in the IMG databases and are linked to the NCBI accession numbers. The 16S rRNA alignment is available upon request from the corresponding author.

## Acknowledgements

The work of THAH and CLN, was funded by the Norwegian Research Council (award 180444/V40). JL and CG were supported by Agence Nationale de la Recherche (ANR) (ANR-12-BSV3-OO23-01). OAP and IVK were supported by the Russian Science Foundation (RSF) grant # 16-14-00121. We thank the Norwegian Sequencing Centre for sequencing and support with the bioinformatics analysis. Strains are available under request at the culture collection UBOCC (www.univ-brest.fr/ubocc) and LM2E, Ifremer collection. We acknowledge Dr. AC Stüken for helpful discussions and feedback on the manuscript.

